# OVGP1 is an oviductal fluid factor essential particularly for early embryonic development in golden hamsters

**DOI:** 10.1101/2022.10.25.513711

**Authors:** Kenji Yamatoya, Masaru Kurosawa, Michiko Hirose, Yoshiki Miura, Hikari Taka, Tomoyuki Nakano, Akiko Hasegawa, Kyosuke Kagami, Hiroshi Yoshitake, Kaoru Goto, Takashi Ueno, Hiroshi Fujiwara, Yoichi Shinkai, Frederick W. K. Kan, Atsuo Ogura, Yoshihiko Araki

**Affiliations:** Institute for Environmental & Gender-specific Medicine, Juntendo University Graduate School of Medicine, Chiba, Japan; RIKEN BioResource Research Center, Ibaraki, Japan; Laboratory of Proteomics & Biomolecular Science, Biomedical Research Core Facilities, Juntendo University Graduate School of Medicine, Tokyo, Japan; Department of Anatomy and Cell Biology, Yamagata University School of Medicine, Yamagata, Japan; Department of Obstetrics & Gynecology, Hyogo Medical University, Hyogo, Japan; Department of Obstetrics & Gynecology, Kanazawa University Graduate School of Medical Sciences, Ishikawa, Japan; Cellular Memory Laboratory, RIKEN Cluster for Pioneering Research, RIKEN, Saitama, Japan; Department of Biomedical and Molecular Sciences, Faculty of Health Sciences, Queen’s University, ON, Canada; Department of Obstetrics & Gynecology, Juntendo University Graduate School of Medicine, Tokyo, Japan; Division of Microbiology and Immunology, Department of Pathology and Microbiology, Nihon University School of Medicine, Tokyo, Japan

**Keywords:** oviductal glycoprotein 1 (OVGP1), knockout-hamster, infertility, early developmental failure, embryonic lethality

## Abstract

The mammalian oviductal lumen is a specialized chamber that provides an environment that strictly regulates fertilization an early embryogenesis, the regulatory mechanisms to gametes/zygote are still largely unknown. In this report, we studied the oviductal regulation of early embryonic development using *Ovgp1* (a gene encoding an oviductal humoral factor, OVGP1)-knockout (KO) hamsters. The experimental results revealed the following: 1) Female *Ovgp1*-KO hamsters fail to produce any litters at all; 2) In the oviducts from KO animal, fertilized eggs are sometimes identified, but their morphology shows abnormal features; 3) The number of implantations in the KO females is evidently low; 4) Even if implantations occur, the embryos develop abnormally and eventually become embryonic lethal; and 5) *Ovgp1*-KO females transferred to wild-type females produce KO egg-derived litters, but the reverse experiment does not. These results suggest that OVGP1-mediated physiological events are crucial for early embryonic development *in vivo*. This animal model shows that the fate of the fertilized egg is not only genetically determined, but that the surrounding oviductal microenvironment plays a pivotal role in normal embryonic development.

**Summary statement:** Deficiency an oviductal humoral factor (OVGP1) caused female infertility in the golden hamsters. The presence or absence of OVGP1 has significant physiological effects on early embryonic development *in vivo*.

## Introduction

The mammalian oviduct is an intra-abdominal organ that serves as the site of fertilization and early embryonic development prior to implantation of the blastocysts in the endometrium. The lumen of the oviduct where fertilization takes place is strictly extracorporeal, as is the lumen of the gastrointestinal tract. Thus, the mammalian reproductive process from fertilization to preimplantation is essentially an *“ex vivo”* event. Because mammalian fertilization appears to take place inside the body, *in vitro* fertilization (IVF), the current treatment method in infertility medicine, is generally misunderstood as a special type of fertilization, but the fact that it takes place outside the body does not make it a special fertilization condition.

The use of culture medium with a clearly defined composition in mammalian IVF was pioneered by Yanagimachi and Chang using the golden hamster (*Mesocricetus auratus*) as an animal model [Yanagimachi & Chang, 1964]. This technique was originally developed to visualize mammalian fertilization. Since then, the theory developed has evolved into an important and fundamental principle for the development of human IVF methods [Edwards et al. 1969]. As a treatment for infertility, especially in humans, IVF-embryo transfer (ET), in which oocytes are harvested directly from the ovaries without passing through the Fallopian tubes (oviducts), fertilized, and cultured in test tubes (dishes), then transferred vaginally into the uterine cavity, has become widely used worldwide. Therefore, oviductal factors are generally considered not always necessarily in the IVF-ET process, and studies of the reproductive physiology of the oviducts have been neglected as a worldwide trend, especially during the last two decades. However, the oviduct (or its homologous organ), the original site of fertilization and early embryonic development, is widely conserved in lower vertebrates as well as in mammals. The origin of sexually reproducing organisms can be traced back to the Cambrian period, at least 600 million years ago [Araki et al, 2021; Araki 2022]. The physiological functions of the oviduct, which have been widely conserved during the long process of evolution and selection, are naturally thought to have important functions (not necessarily one) which are not yet known.

The medium for IVF-ET has a long history of development based on the composition of the original Fallopian tube and uterine fluids [Quinn et al, 1985ab; Gardner et al, 1996]. The composition was based on the concentrations of glucose, inorganic salts, growth factors and hormones, but at that time the proteins, which are the main components of oviductal fluid, were largely unknown. Therefore, serum components were used as a substitute for the other components of the oviductal fluid. However, the ultimate goal of IVF should be to reproduce the *in vivo* oviductal microenvironment as closely as possible, as a medical treatment. It is also biologically important to elucidate the reproductive physiology of the oviduct, or the site of fertilization and early embryonic development in the majority of mammals.

In this study, using the golden hamster as a prototype model for mammalian IVF, we have provided evidence to demonstrate that a humoral factor in the oviduct, Oviductal glycoprotein 1 (OVGP1) has a significant effect on early embryonic development *in vivo*.

## Results

### Generation of OVGP1-deficient hamsters

Among oviductal factors identified in the oviductal fluid, OVGP1 has been well characterized and suggested to play important roles in the process of several mammalian species, including humans [for review, see Araki & Yoshida-Komiya, 1998; Buhi 2002; Avilés et al, 2010; González-Brusi et al, 2020; Zhao et al, 2022]. Using the golden hamster as a model, which has provided a wealth of knowledge concerning the mammalian reproductive process [Hirose & Ogura, 2019], *Ovgp1*-knockout (KO) animals were generated using gene editing technology (Suppl. Fig. 1A,B). Preliminary mating experiments showed that the F0 KO females (2 independent individuals; #8, #10) did not produce any offspring. On the other hand, KO males (#3, #4) were found to be as fertile as those of wild-type (WT). In the F1 and later generations, fertility was confirmed in 17 of 23 (73%) KO male individuals. Fertility was also confirmed to be possible in heterozygous females; 28 of 45 (62%) heterozygous-female individuals were confirmed fertile with an average litter size of 7.08, in the post-F1 generation. Therefore, we maintained a line of KO males and heterozygous females for further experiments.

Western blotting with OVGP1-specific antibodies did not reveal OVGP1 expression signal in *Ovgp1*-KO hamster oviducts (Suppl. Fig. 1C). In additional mating experiments with F0-F2 KO females with WT males with confirmed fertility (total pair number: 15), no litters were obtained from all *Ovgp1*-KO females (Suppl. Fig. 2A). These results suggest a lack of fertility in *Ovgp1* KO females. However, a F0 female (#10) used in the mating experiment did not show any outward signs of pregnancy, but went into shock and died suddenly at 15-day-post-coitus (dpc)(almost full-term) after several mating sessions (Suppl. Fig.2B). An autopsy revealed that the death was almost sudden, as there was a large amount of food in the gastrointestinal tract. Both uterine horns were externally hematomatous, and the split surface had a hematoma visible to the naked eye around the fetal sac, but no fetus was observed (Suppl. Fig. 2Ba; 2Bb). After fixation, observation under a light microscope revealed a placenta-like structure, but no fetal scar was noticeable due to absorption and hemorrhage (Suppl. Fig. 2Bc). Since this phenomenon was limited to this one case and was an F0 individual, we cannot conclude at this time that it was the result of *Ovgp1* gene editing. However, this was considered a clear example of how OVGP1 may have a significant effect on the reproductive process in the hamster models.

### Fertilizing ability of *Ovgp1*-KO females

In order to find out what happened during the reproductive process, we first went on to examine the development of the eggs after mating. In *Ovgp1*-KO animals, some of the 1-dpc eggs appeared to be fertilized, but not most as in WT (Fig. 1A (control), D). When examined under the light microscope (binary image), the egg cytoplasm from the KO hamsters showed a central accumulation of intracellular organelles (Fig. 1B (control), E). Electron microscopy showed that the KO eggs at 1-dpc displayed a thinner zona pellucida (ZP) and a heterogeneous distribution of intracellular organelles (Fig. 1C (control), F). In contrast to the synchronous development of four to eight cells in fertilized eggs in WT 2.5-dpc, eggs in the oviducts of *Ovgp1*-KO females showed developmental abnormalities evident at the level of light microscopy, such as delayed development, disproportionate egg breakage and degeneration (Suppl.Fig.3).

**Figure 1.**
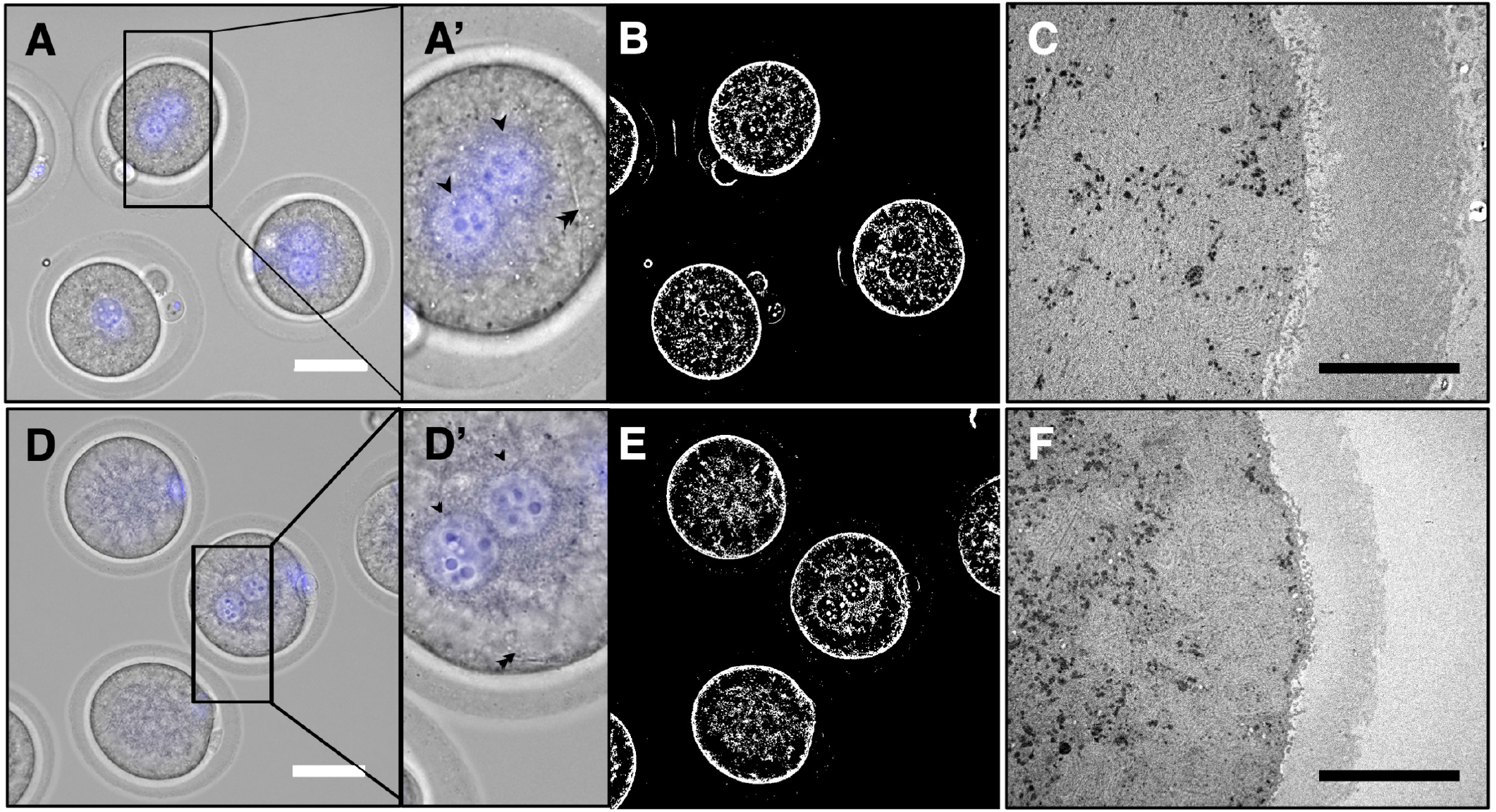
Morphological findings of zygotes at 1-dpc in hamsters. Female animals were mated spontaneously with fertile WT males. Eggs were collected from the ampullary region of oviduct, and counter-stained with 4’,6-diamidino-2-phenylindole solution (FUJI FILM Wako Chemicals) for light microscopic (LM) observation and some were processed for electron microscopy (EM). The images show eggs from WT female (A, B; binary image of A, C; transmission electron microscopic (TEM) image) and *Ovgp1*-KO female (D, E; binary image of D, F; TEM image), respectively. Pronuclei indicated by arrowheads and sperm tails are shown by double arrowheads in the high magnification images (A’, D’). Bars; 50 μm (LM); 10 μm (EM)

At 4.5-dpc, implantation sites were observed in WT animals but not in the KO animals (Fig. 2A). At the time of this experiment, it was thought that early embryos might not implant in *Ovgp1*-KO female animals. However, at 5.5-dpc, statistically significant fewer implantation sites were observed in the KO hamsters (Fig. 2B). These results suggest that at least a small number of embryos produced by the KO females were able to implant in the uterus.

**Figure 2.**
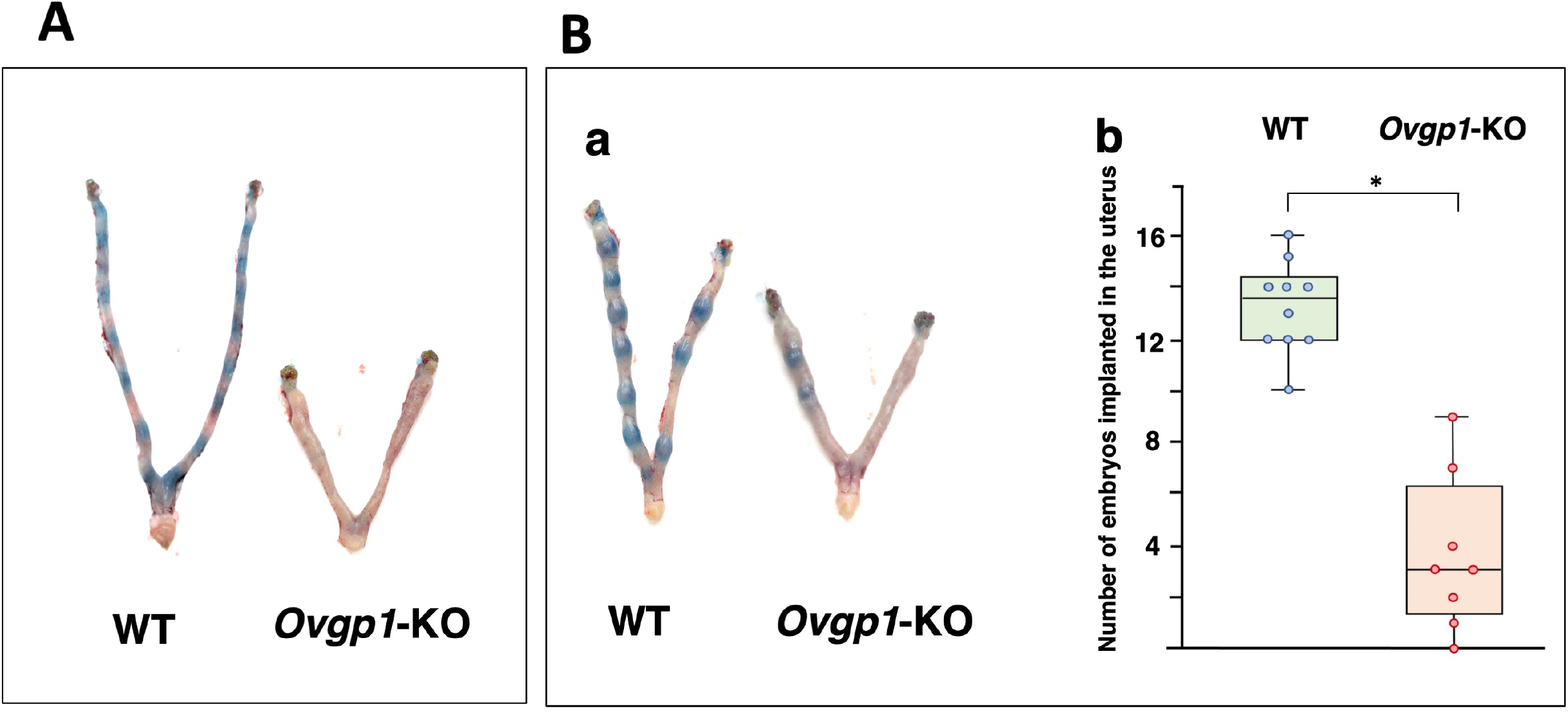
Observation of fetal implantation in the uterus of hamsters. At 4.5-dpc (A), and 5.5-dpc (B). Female animals were mated spontaneously with fertile WT males, and showed typical uterine appearance at 4.5-dpc and 5.5-dpc (b). Box-and-whisker diagram of the number of implantations at 5.5-dpc (c). *P*-value was calculated by the Mann-Whitney *U* test. * *P*<0.05

### Histological findings of 8.5-dpc embryos of *Ovgp1*-KO females

At 8.5-dpc, WT implanting embryos were well developed and pregnancy was clearly visible (Fig. 3A-a). On the contrary, in *Ovgp1*-KO females, the number of implanting embryos at 7.5-8.5-dpc was small or not well visible to the naked eye (n=4). Among them, one individual appeared to be as well developed in appearance as those of the WT (Fig, 3A-b).

**Figure 3.**
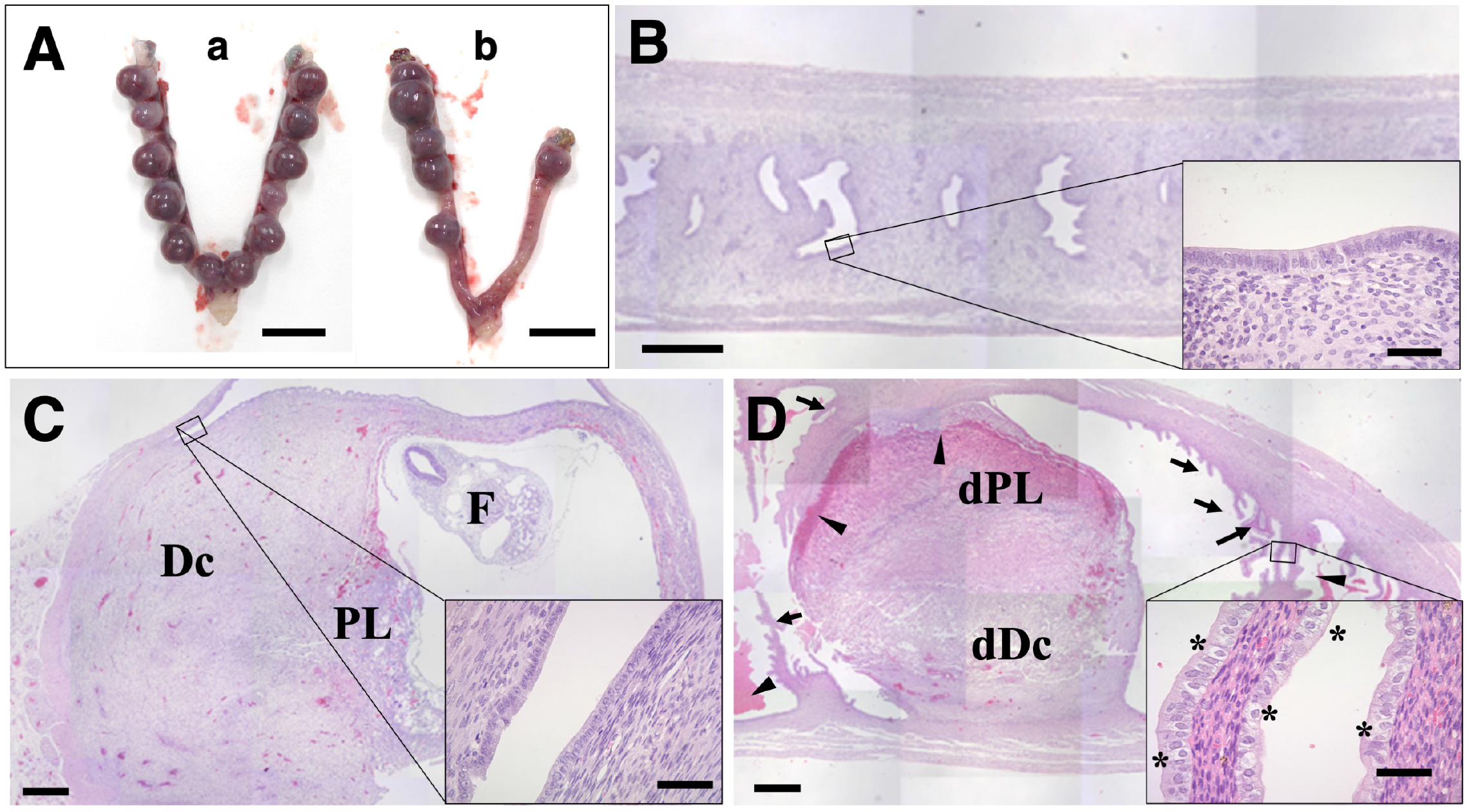
Morphological findings of the uteri at 8.5-dpc in hamsters. (A) Appearance of the uteri in pregnancy; WT (a) and *Ovgp1*-KO (b). Bars: 10 mm. (B) Sagittal section of the uterus of a KO hamster in non-pregnant state. The inset shows a magnified view of the endometrium. Bar: 500 μm (inset: 50 μm). Sagittal sections of pregnant uteri and their corresponding magnified images (insets) of WT (C); *Ovgp1*-KO (D). Dc: decidua cells; PL, placenta; F, fetus. dPL; hemorrhagic degenerated placental tissues, dDC; denatured decidualized membrane cells. Arrowheads indicate hemorrhage traces, and arrows reveal endometrial folds not seen in WT. Cuboidal epithelial cells with spherical nuclei and bright cytoplasm are shown by asterisks (inset). Bars: 500 μm (50 μm in insets).

The uterine epithelium of *Ovgp1*-KO hamsters (non-pregnant) showed no obvious abnormality under the light microscope (Fig. 3B) when compared to the uterine epithelium of WT hamsters. At 8.5-dpc of WT, the embryo was well developed and the developing embryo and placenta can be seen in the fetal sac (Fig.3C). The developing embryo (fetus) can be seen with the naked eye. The endometrium, except for the implantation sites, can be seen to be composed of single columnar epithelium (Fig. 3C, inset).

In *Ovgp1*-KO hamsters, no developing fetus was observed either with the naked eye or under the stereomicroscope. When examined under a light microscope, placenta/decidua-like primordial tissue with hemorrhagic degeneration was observed, but no trace of embryo buds could be seen, and there was marked hemorrhage in the placenta and uterine cavity (Fig.3D, arrowheads). It should be noted that the endometrium adjacent to the implantation site differed from that of the WT in that it showed formation of epithelial folds reminiscent of the ampulla of the oviduct (Fig.3D, arrows). High magnification of this area revealed the presence of numerous cuboidal cells each with a spherical nucleus and pale cytoplasm (Fig.3D, inset, noted by asterisks). These are probably secretory cells intercalating with non-secretory cells reflective of the typical columnar epithelium found in the WT.

### Validation of phenotypic recovery after ovarian transplantation

Female hamsters in which the *Ovgp1* coding region was inactivated by gene editing became completely infertile. This is strongly suggested to be the result of the absence of OVGP1 in the oviductal microenvironment of the KO female hamsters, which adversely affects early embryonic development immediately after fertilization. However, in order to confirm that this clear phenotype is correct, it is usually necessary to knock-in the inactivated gene region and see if the phenotype is restored. At present, however, there are technical limitations to achieve this in hamsters, since they have more fragile eggs than mice. Instead, we tried to see if we could restore the phenotype using the ovarian transplant technique (Fig.4).

**Figure 4.**
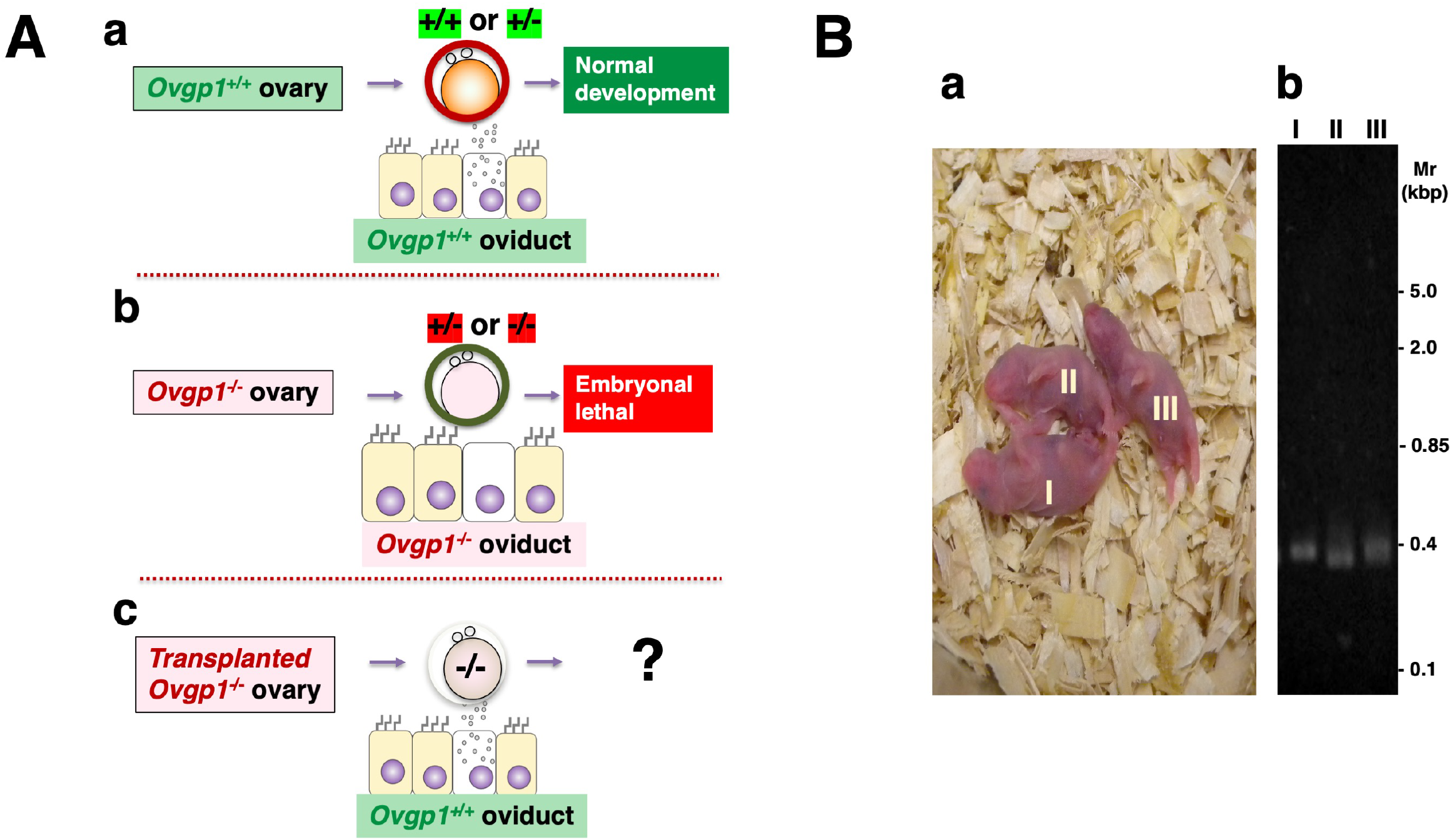
Effects of OVGP1 on early embryogenesis. (A) Schematic of fertilized eggs and oviductal epithelium in early development: Post-fertilization embryos and oviductal epithelium in WT (a) and *Ovgp1*-KO (b) females mated with either WT, heterozygous (*Ovgp1*^+/-^) or KO male, and what happens if a WT female is implanted with an *Ovgp1-*KO ovary and mated with an *Ovgp1*-KO male?(c). (B) Typical results of ovarian transplantation experiments proposed in A-c showing the appearance of litters (a) and their genotyping (b).

The fact that *Ovgp1*-KO male hamsters are fertile as described above, indicates that even if the genotype of the zygote is *Ovgp1*^+/-^, the embryo can develop normally and produce litters in the wild-type oviduct microenvironment (Fig.4Aa). On the other hand, embryos in the *Ovgp1*-KO female oviducts are lethally altered and no viable offspring are produced whether the eggs are genotyped *Ovgp1*^+/-^ or ^-/-^ (Fig.4Ab).

What happened when these WT females were implanted with KO ovaries and mated with KO males (Fig. 4Ac)? As expected, these experiments resulted in *Ovgp1*^-/-^ litters (out of a total of 26 transplant experiments, litters were obtained from 11 individuals) that could never be obtained by normal mating (Supple. Table S1; Fig.4B). These litters can only be obtained under natural conditions by mating *Ovgp1*^-/-^ males with *Ovgp1*^+/-^ females. Conversely, transplantation of the ovaries from WT individuals into KO females did not result in any offspring (n=5): it is noteworthy to mention that preliminary experiments of ovarian transplantation using WT females as both recipients and donors, showed that 11 out of 14 (78.6%) transplanted animals produced litters. The possibility that an individual transplanted with ovaries from a WT individual to a KO female could produce litters cannot be ruled out. However, when ovarian transplants were performed using a 78.6% success rate technique, no litters were produced from all five individuals. The probability of such an event occurring can be calculated to be less than 1%.

### What’s going on in the *Ovgp1*-KO hamster oviduct: comprehensive quantitative protein analysis in the oviduct after ovulation

As noted above, OVGP1 is strongly suggested to be a microenvironmental factor in the hamster oviduct, the site of fertilization and early embryonic development, with pronounced effects on gametes and zygotes.

At this time, it is unclear whether the results obtained in the present study regarding the physiological activity of OVGP1 is commonly found in other mammals. However, at the very least, exploring what is happening in the hamster oviduct represents an important first step toward understanding the molecular mechanisms of these phenomena. Therefore, an attempt was first made to perform a comprehensive protein microanalysis by confining the target especially to the oviduct with unfertilized eggs retained immediately after ovulation in their lumen prior to mating.

Quantitative data for a total of 3,572 oviductal proteins were obtained by mass spectrometry (MS) of the oviducts of super-ovulated animals (3 WT and 3 *Ovgp1*-KO, each containing an unfertilized egg-cumulus complex in the oviduct lumen) (MS data have been deposited in ProteomeXchange and jPOST with the accession codes PXD037067 and JPST001867, respectively). In the Volcano diagram, changes in the expression of oviductal proteins associated with OVGP1 deficiency were observed. Of the 21 proteins that showed significant expression variation, a relatively large number of down-regulated proteins (18 proteins) were identified, including some the function of which has been previously implicated in the reproductive processes (Fig. 5; Supple. Table S2).

**Figure 5.**
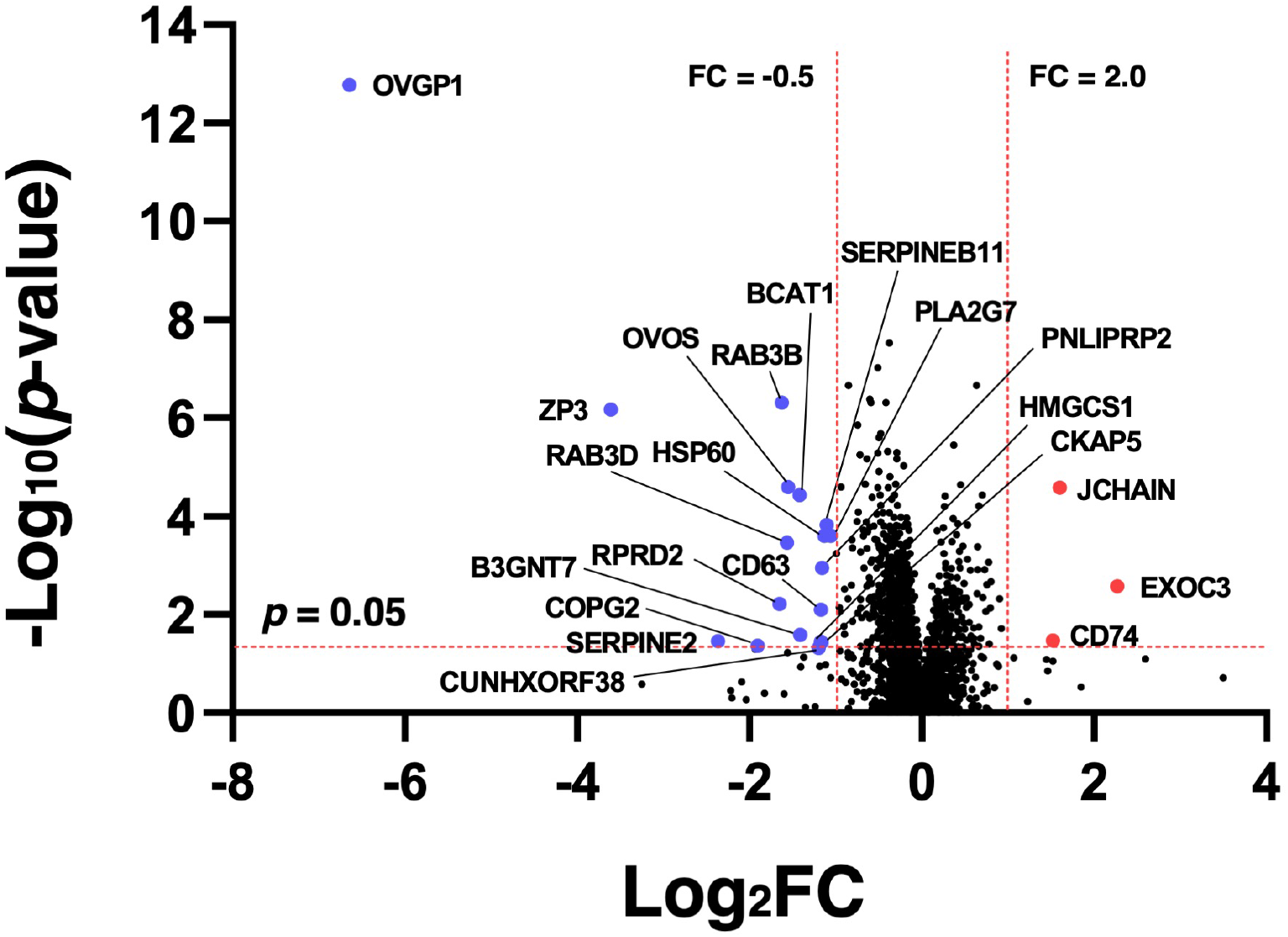
Volcano diagram showing the down/up-regulated oviductal proteins after ovulation in WT/*Ovgp1*-KO females. Oviducts were isolated from each of three independent WT/*Ovgp1* KO individuals. Dotted red lines indicate the fold-change (FC) protein expression at 2 and −0.5, and *p*-value = 0.05, respectively. Blue/red dots show the down/up-regulated proteins in the oviducts from *Ovgp1*-KO females. BCAT1; branched-chain-amino-acid aminotransferase, B3GNT7; hexosyltransferase, CCDC114; coiled-coil domain-containing protein 114, CD63; tetraspanin, CD74; HLA class II histocompatibility antigen gamma chain isoform X2, CKAP5; cytoskeleton-associated protein 5, COPG2; Coatomer subunit gamma-2, CUNHXORF38; uncharacterized protein CXorf38 homolog isoform X1; EXOC3; exocyst complex component 3, HMGCS1; hydroxymethylglutaryl-CoA synthase, HSP60; 60 kDa heat shock protein, mitochondrial, JCHAIN; immunoglobulin J chain, OVGP1; oviduct-specific glycoprotein 1, OVOS; ovostatin homolog, PLA2G7; platelet-activating factor acetylhydrolase, PNLIPRP2; triacylglycerol lipase, SERPIN2; glia-derived nexin (serpin family E member 2), SERPINB11; serpin family B member 11, RAB3B/D; Ras-related protein Rab-3B/D, RPRD2; regulation of nuclear pre-mRNA domain containing 2, ZP3; zona pellucida sperm-binding protein 3. KRT90; keratin, type II cytoskeletal cochlear-like, is considered a contaminant in sample preparation and is listed in Supplementary Table S2 but omitted in this figure.

## Discussion

This study has clearly shown, for the first time, that a deficiency of a fluid factor secreted into the lumen of the oviduct causes lethal changes in early embryonic development. Results of the present study are almost consistent with the currently proposed bioactivities of OVGP1, which has been demonstrated in a variety of mammalian models to date, including humans [Araki & Yoshida-Komiya, 1998; Buhi 2002; Avilés et al, 2010; González-Brusi et al, 2020; Zao et al, 2022] summarized in Figure 6. Although OVGP1 has been suggested to be involved in sperm functions, fertilization, and embryonic development, the clear KO phenotype observed in the present study is a lethal change in embryonic development after fertilization. Since normal embryonic development involves a comprehensive process including sperm-egg function and fertilization, therefore, it is premature to conclude from this phenotype that OVGP is essential only for embryonic development.

**Figure 6.**
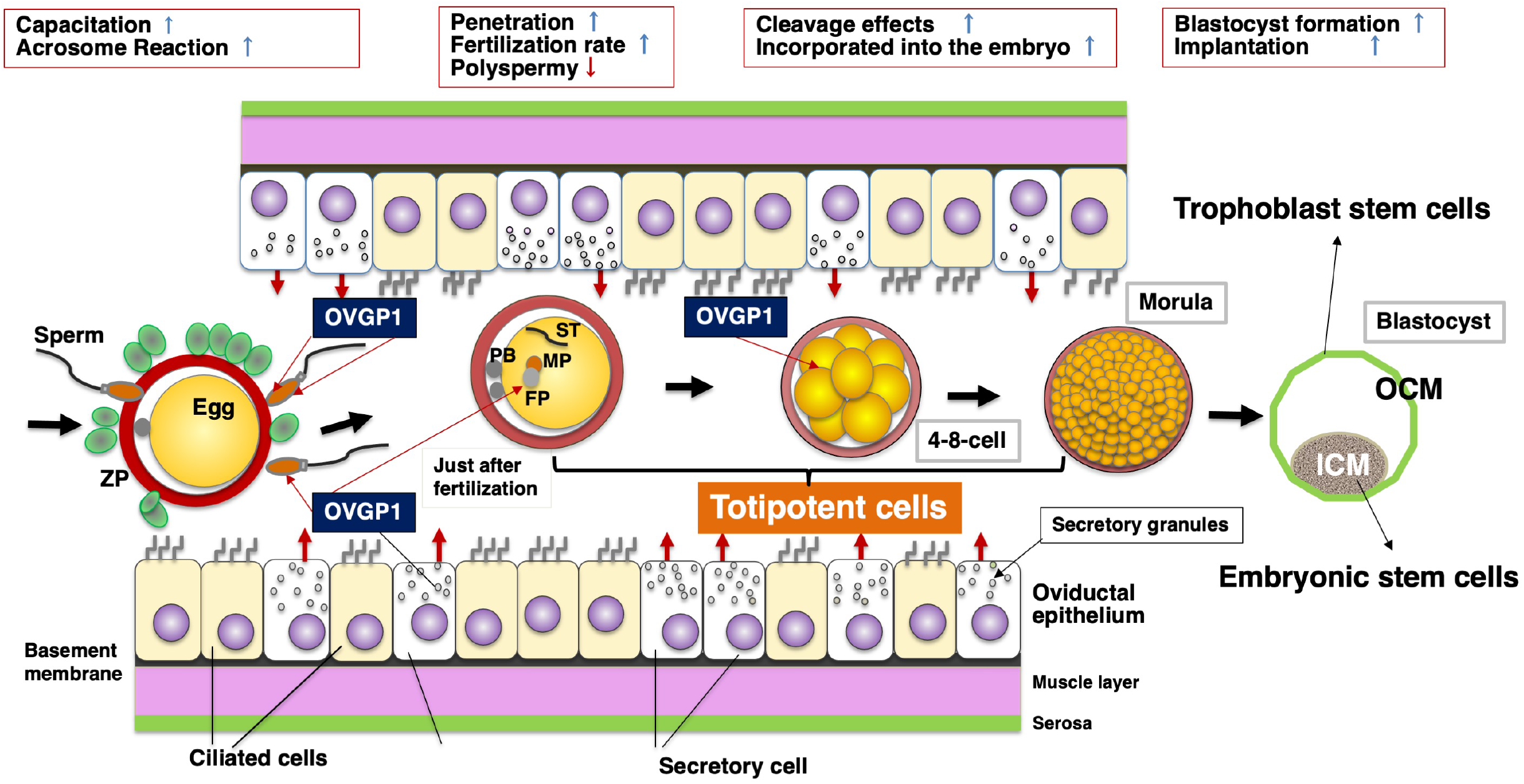
Potential physiological activities of an oviductal humoral factor OVGP1 during early reproductive process as suggested by the present study. Previous *in vitro* experiments have proposed that OVGP1 secreted from the oviduct 1) modifies the egg, 2) also modifies the sperm, and 3) is incorporated into the early embryo. The present study has demonstrated that OVGP1 is an essential factor that aids fertilization and the normal development of early embryos (totipotent cells) in the hamster model. ZP, zona pellucida; PB, polar body; MP, male pronucleus; FP, female pronucleus; ST, sperm tail; ICM, inner cell mass; OCT, outer cell mass.

In hamsters, OVGP1 has been suggested to be an important fluid factor during *in vivo* fertilization, especially since it modifies the ZP of oocytes in transit in the oviduct after ovulation [Araki et al, 1987; Oikawa et al, 1988; Robitaille et al, 1988] and inhibition of IVF was shown by targeting OVGP1 with a specific antibody [Sakai et al, 1988]. However, the conditions of experiments *in vitro* carried out with OVGP1 are very different from their counterparts carried out *in vivo*, so that it is natural to argue that we should be more cautious about evaluating *in vitro* interpretations as compared to *in vivo* functions [O’Day-Bowman et al, 2002]. In addition, OVGP1 has been found to be taken up by the developing embryos after fertilizations [Kan et al, 1993], suggesting that it may have a role in early embryonic development and may even have a function in implantation [Roux et al, 1997]. The data obtained in the present study provide direct evidence of the intrinsic importance of the physiological function of OVGP1 in the reproductive process of the hamster.

In considering the results of this study, it is necessary to reconsider the implications of the lack of a clear phenotype in *Ovgp1*-KO mice published 20 years ago (these genetically modified animals produced litters comparable to WT) [Araki et al, 2003]. At present, we cannot elaborate on the phenotypic differences between *Ovgp1*-KO mice and hamsters due to insufficient experimental data. However, it is known that the primary structure of OVGP1 is highly conserved between species on the N-terminal end, and that there is considerable structural diversity between species on the C-terminal end and in its degree of glycosylation [Araki & Yoshida-Komiya, 1998; Zao et al, 2022]. Furthermore, until the end of the 20th century, the molecular characterization of OVGP1 was mainly carried out in large livestock such as cattle, pigs and sheep where relatively large amounts of oviductal fluid were available, and in baboons as a human model. Previous studies on the functions of OVGP1 in the reproductive process have been carried out in rodents using mainly hamsters, but not mice, because of the stability of their sexual cycle. As for mouse OVGP1, there were only three reports all from the same research group in the mid-1980s [Kapur & Johnson, 1985; 1986; 1988], and there have been no further protein-level reports to date concerning mouse OVGP1. Since the identification of OVGP1, knowledge about genetic engineering of mouse oviductal glycoprotein has been accumulating [Sendai et al, 1995; Takahashi et al, 2000], while studies of the underlying mechanisms of OVGP1 that regulate its function are clearly lacking. This is one of the reasons why research in this field has not progressed since the establishment of the *Ovgp1*-KO mouse.

Elucidating the molecular mechanisms of the phenotype caused by OVGP1 deficiency is an urgent research priority. For this reason, the results of MS analysis using oviducts immediately after ovulation (Fig. 5; Suppl. Table 2) could be one of the breakthroughs. Among the group of molecules in the oviduct down-regulated by *Ovgp1* KO hamsters, ZP3, the major component of ZP glycoproteins, deserves the first attention. Although the importance of the ZP glycans and the structure and function of homologous molecules among all animal models examined to date remains controversial [Tulsiani et al, 1988; Moros-Nicolás et al, 2019; Tumova et al, 2021], ZP3 has been suggested to play an important role in fertilization due to the protein’s three-dimensional structure [Tracy et al, 1995; Han et al, 2010]. Similarly, serpin family E member 2 (SERPIN2) and ovostatin homolog (OVOS), serine protease inhibitors, were identified as down-regulated molecules by *Ovgp1*-KO hamsters (Fig. 5). SERPIN2 is abundantly expressed in granulosa cells although its function is unknown in humans [THE HUMAN PROTEIN ATRAS; https://www.proteinatlas.org/ENSG00000135919-SERPINE2]. Furthermore, OVOS has also been identified in mouse uterine fluid [Huang et al, 2019] and may also be present in oviductal fluid. These findings suggest that since unfertilized eggs are surrounded by cumulus cells (*i.e*., granulosa cells) in the oviductal lumen, the decrease in SERPIN2 and OVOS may have caused a local increase in protease activity, which in turn, affected ZP3 stability and decreased fertilization rates. In hamsters, OVGP1 modifies the ZP promptly after ovulation [Araki et al, 1987; Oikawa et al, 1988; Robitaille et al, 1988], and its loss may directly or indirectly affect the three-dimensional structure of the ZP by downregulating ZP3. The latter speculation is also consistent with the thinning of the ZP immediately after fertilization as shown by electron microscopy (Fig. 1F). In general, the current status of research on hamster OVGP1 is that various databases are still underdeveloped compared to those of humans and mice. However, since this is a problem that will be resolved over time, we believe that the present study has provided sufficient and novel results that could considered as a breakthrough in OVGP1 research.

Early mammalian development (even before the widespread use of recombinant technology) has been studied mainly in the mouse model. This is because the development of blastocysts is easier in media with a simpler chemical composition, but this development process is much more complex in the *in vivo* situation in mammals. *In vitro* culture of non-mouse embryos, including humans, still has some technical issues to overcome such as two-cell block and blastocyst formation, and it took a long time to solve these problems in hamsters through various innovations [Schini & Bavister, 1988]. Although it is now possible to culture hamsters to blastocysts and obtain litters by embryo transfer [Seshagiri & Vani, 2019], the unfertilized eggs used in these experiments, whether in hamsters or mice, are usually prepared from oviducts after ovulation. These facts suggest that the microenvironment in the oviduct and uterus, which is generally important for early embryonic development, is governed by a variety of cell biological control mechanisms. The *in vivo* interaction between the early embryos and the epithelium of the female reproductive tract can be inferred from the fact that early embryos, which are undifferentiated totipotent cells, have a much higher mitotic potential than cancer cells (it is almost unlikely that a single cancer cell will grow to the same weight as fetal tissue in the same amount of time as its gestation period, not to mention the human example), and are reliably regulated to carry out normal development.

At present, IVF-ET is widely used as a treatment for infertility in which eggs from the ovaries are fertilized in a culture medium and trans-vaginally transferred into the uterine cavity for subsequent implantation, but the reproductive physiology of the oviducts has been long neglected as described above. However, the oviducts (and their homologous organs), the original site of fertilization and early embryonic development, are widely conserved in vertebrates as well as mammals, and one could imagine that they have important physiological functions yet to be explored and unraveled. The molecular comparison of the reproductive processes between the *Ovgp1*-KO hamsters established in the present study and the homologous gene KO mice provides an excellent animal model to further elucidate the mammalian reproductive mechanisms.

Comprehensive molecular dynamics studies are currently underway using these KO animals. Future research originating from OVGP1 has the potential to elucidate some aspects of the pathogenesis for human disease, such as infertility due to disorders of the early fertilization process or infertility due to fetal growth retardation in the early stage of pregnancy the causes of which remain unknown.

## Materials and Methods

### Animals

Sexually mature (7~8-week-old) golden hamsters (*Mesocricetus auratus*) were purchased from Japan SLC, Inc. (Hamamatsu, Shizuoka, Japan). They were maintained and bred at our Animal facilities under 12L:12D conditions. Observation of sperms in the vaginal plug in females after mating was considered as 0-dpc. All animal experiments were conducted according to the guidelines for care and use of laboratory animals, Juntendo University (approval # 768) and RIKEN Tsukuba Institute, (approval # T2021-Jitsu004) Japan.

For genotyping, ear biopsies were lysed with 0.4 mg/mL proteinase K (Nakalai Tesque Inc., Kyoto, Japan) and partially purified using standard chloroform extraction. Genomic fragments containing the target site were then amplified by PCR using primers (forward: 5’-AAGCCAGAATCCAAAGCTGAAGCAC-3’; Reverse: 5’-GTATTAAACCCTCACAACTGGGCTC-3’). The PCR procedure followed the instructions for Tks Gflex DNA polymerase (TaKaRa Bio Inc., Shiga, Japan). The amplified PCR fragments were subcloned into the pGEM T Vector system (Promega Corporation, Madison, WI, USA) and sequenced to confirm each allele.

### Generation of *Ovgp1*-KO hamsters

*Ovgp1*-KO hamsters were established using an *in vivo* electroporation CRISPR–Cas9 system, essentially followed as described previously [Hirose et al, 2020]. Pairs of sgRNAs were designed to delete the *OVGP1* genomic sequence from exon 1 to 3 (sequence of DNA targets: + allele; 5’-ACTGACTCCCTGCTAGCGTCAGG-3’, - allele; 5’-CCTGCTAGCGTCAGGCCACGGAT-3’; 5’-CCATCGACCAGCCCCCTGAGCTG-3’; 5’-CCTCGATGACTTGGGAGTTAATG-3’)(Suppl. Fig.1A). Ten animals were born of which three male individuals (#1, 3, 4) and two females individuals (#8, 10) appeared to be homozygously defective in the target gene region (Suppl. Fig. 1B). Females #8 and #10 did not show any external signs of pregnancy in mating experiments with wild-type males with confirmed fertility; two males (#3, 4) were fertile and the defective gene was transmitted to their offspring when mated with wild-type females. To minimize the possible effects of off-targeting when heterozygotes were mated, two generations of heterozygotes were mated to wild-type and the heterozygotes were mated to homozygous males derived from #3 and #4, respectively, and were found to produce a normal number of offspring.

### Western blotting analysis

The concentration of total protein extracted from animal organs was quantified using BradfordUltra (Expedeon Ltd, Cambridgeshire, UK). Antibodies (Abs) specific for hamster OVGP1 used in this study were as follows: AZPO-8 (monoclonal Ab (mAb) against the oligosaccharide portion of hamster OVGP1 (mouse IgG1)[Araki et al, 1987]); anti-OVGP1 N-terminal -peptide polyclonal Ab (pAb)(rabbit IgG, ab74544; Abcam plc, Cambridge, UK); horse radish peroxidase (HRP)-conjugated anti-mouse IgG pAb (P0260) and anti-rabbit IgG pAb (P0448)(Dako, Carpinteria, CA, USA). Proteins were separated by SDS-PAGE system and transferred to Immobilon-P membrane (Merck KGaA, Darmstadt, Germany). Immunoreactions were detected according to standard methods described previously [Yoshitake et al, 2015; Oda-Sakurai et al, 2019].

### Collection of eggs

Eggs were collected from the oviducts of mature females by natural mating with fertile males. Coitus was confirmed by the presence of vaginal sperm in the post-ovulatory vaginal discharge and that day was defined as 0-dpc. For the collection of unfertilized eggs in the oviduct, ovulation was artificially induced by gonadotropin according to the standard method as described previously [Araki et al, 1987; 1992]

### Morphological observation

Tissues from animals were fixed with 20% Formalin solution (FUJUFILM Wako Chemical Co., Osaka, Japan) and embedded in paraffin wax according to standard procedure. Three-μm thick sections were cut and stained with Hematoxylin-Eosin for light microscopic observations. For transmission electron microscopy, samples were fixed with 2.5% glutaraldehyde (FUJIFILM Wako Pure Chemical) in 0.1 M phosphate buffer (PB)(pH 7.2) followed by post-fixation with 2% OsO_4_ in 0.1 M PB (pH 7.4). Fixed specimens were dehydrated with a graded series of ethanol, and embedded in Epok812 (Okenshoji Co., Ltd. Tokyo, Japan) according to standard procedure.

To detect implantation site(s), a solution of Chicago Sky Blue 6B (Tokyo Kasei Kogyo Co., Ltd.) diluted to 1% in saline was injected into the heart of female animals under anesthesia. After circulating for 10 min, blood was perfused with 50 mL of PB saline to clearly visualize the blue pigment in the uterus.

### Ovary transplantation

The technique of ovarian transplantation used in this study essentially followed the methods reported elsewhere [Yun et al, 1990; Takahashi et al., 2001; Miyoshi et al, 2002].

### Statistical analysis

To compare the implantation number at 5.5-dpc of WT and *Ovgp1* KO female animals, the Mann-Whitney *U* test was utilized. A probability of *p* < 0.05 was considered statistically significant.

### Quantitative protein MS analysis

#### a) Sample preparation and protein digestion

Three oviducts collected at after ovulation induced by hormonal induction from each of WT and *Ovgp1*-KO hamster were used for analysis. Samples were lyophilized and stored at −80°C until use. Protein digestion on S-Trap™micro (ProtiFi, Huntington, NY, USA was performed according to the manufacturer’s procedure, except for reductive alkylation. Briefly, lyophilized samples were mixed with lysis buffer containing 5% SDS, 4 mM tris (2-carboxyethyl) phosphine, 16 mM chloroacetamide, 50 mM triethylammonium bicarbonate (TEAB)(Thermo Fisher Scientific Inc., Waltham, MA, USA) boiled at 95°C for 5 min, and cooled to room temperature for 30 min. Afterward, phosphoric acid was added to oviductal lysate to a final concentration of 1.2%, and then six volumes of binding buffer containing 90% methanol, 100 mM TEAB were added. The protein solutions were loaded to an S-Trap filter, spun at 4,000 × g for 30 sec. Then the filter was washed 3 times with 150 μL of binding buffer. Finally, 2 μg of MS grade Trypsin Platinum (Promega, Madison, WI, USA) in 40 μL of digestion buffer containing 50 mM TEAB was added into the filter and digested at 37°C for 16 h. To elute peptides, three stepwise buffers were applied, with 40 μL each containing 50 mM TEAB, 0.2% formic acid in water, and 50% acetonitrile. The peptide solutions were pooled, lyophilized, and desalted with a GL-Tip SDB column (GL Sciences, Tokyo, Japan).

#### b) Liquid chromatography (LC)/MS analysis

LC/MS analysis was performed using an Orbitrap Eclipse Tribrid mass spectrometer equipped with FAIMS PRO coupled to an EASY-nLC 1200 system (Thermo Fisher Scientific Inc.). Peptides were resuspended in a mixture of 0.1% formic acid and 2% acetonitrile (v/v), and then loaded on a packed C18 column (15 cm, 3 μm, 75 μm i.d., Nikyo Technos Co.,Ltd, Tokyo, Japan). As mobile phases, 0.1% formic acid in water was used for mobile phase A and a mixture of 0.1% formic acid and 90% acetonitrile (v/v) for mobile phase B. Peptides were separated with a linear gradient of acetonitrile from 2% to 35% (phase B) at 120 min at flow rate of 300 nL/min. The mass spectrometer was operated in positive ionization mode, and the correction voltages on the FAIMS Pro interface were set to −40V, −60V, and −80V, respectively. Data-dependent acquisition (DDA) was performed using the following parameters. MS1 resolution was set at 60,000, and a maximum inject time was set to auto and scan range from 350 to 1,800 *m/z*. During the tandem MS (MS/MS) scan, the linear ion trap analyzer detects, and the precursor sets its intensity threshold at 5,000. For precursor fragmentation in High-energy collisional dissociation (HCD) mode, a normalized collision energy of 30% was used.

Protein identification and label-free quantification were performed using Proteome Discoverer (PD)(ver. 2.5.0.400, Thermo Fisher Scientific Inc.). SEQUEST HT in PD and MASCOT (Version 2.8.0, Matrix Science Inc.) were used as search engines, and all raw files were searched against a *Mesocricetus auratus* (Syrian Golden hamster) protein database in Universal Proteins Resource Knowledgebase (UniProtKB)(32,230 entries, 2022_02). Carbamidomethylation of cysteine was a fixed modification, while acetylation of the protein N-terminus and oxidation of methionine were variable modifications. Mass tolerances were set at 10 ppm and 0.8 Da for MS and MS/MS, respectively. A false discovery rate (FDR) of 1% was applied to the analysis for both peptide and peptide spectral match levels. Peaks were detected and integrated using the Minora algorithm embedded in PD. Proteins were quantified based on unique and razor peptides intensities. Normalization was performed based on the total protein amount.

## Acknowledgments

We are indebted to Dr. Satoshi Hayakawa (Nihon University), Dr. Ichiro Miyoshi, Dr. Shoichiro Kurata (Tohoku University), and Dr. Hiromi Yoshida-Komiya (Fukushima Medical University) for both material and emotional support. The authors express deep appreciations to Dr. Soichiro Kakuta (Juntendo University) and Dr. Kei Fukuda (RIKEN) for their technical supports in using electron microscopy and designing gRNAs, respectively. The excellent secretarial assistance of Ms. Keiko Fukuda (Jundendo University) and Ms. Mari Enomoto (Nihon University) is gratefully acknowledged.

## Data availability

Data for comprehensive protein quantitative analysis by mass spectrometry reported in this paper have been submitted to the ProteomeXchange and jPOST with the accession codes PXD037067 and JPST001867, respectively.

## Figure legends

**Supplementary Figure 1**. Production of *Ovgp1*-null hamsters. Gene structure of hamster *Ovgp1* and its editing strategy (A). The position of the gene sequence to be removed from EXON 1 to 3 is indicated by vertical arrows and the position of PCR primers for mutant detection is indicated by red arrows. Genotypes of F0 animals after gene editing by PCR (B). The positions of the predicted PCR products relative to the genomic DNA are indicated by arrowheads (WT) and double arrowheads (KO), respectively. Male #1, 3, 4 and female #8, 10 were successfully gene edited as designed. Male #1, females #8, and #10 did not produce pups, so males #3 and #4 were used to maintain the strain. #4(F3) indicates that DNA extracted from a F3 generation female derived from a #4 F0 male individual was used as the template. Western blotting analysis using OVGP1-specific antibodies (C); Equal amounts of tissue protein solution were detected by SDS-PAGE followed by OVGP1-specific antibodies (AZPO8, recognizing carbohydrate moiety of the OVGP1 (a); ab74544, recognizing the N-terminal peptide of OVGP1 (b)). lanes 1: ovary, 2: oviduct and 3: uterus, respectively.

**Supplementary Figure 2**. Reproductive ability of *Ovgp1*-KO hamsters. Fertility of *Ovgp1*-KO female hamsters (A). Female WT (n=5) and *Ovgp1*-KO (n=15) mated with fertility confirmed WT males. Autopsy image (B) of an F0 *Ovgp1*-KO female (15-dpc) that died suddenly during a mating experiment. Appearance of the uterus (a), its cross-sectional image (b) and hematoxylin-eosin stained image (c). Bar = 1 mm.

**Supplementary Figure 3**. Early embryos at 2.5-dpc. From oviduct of WT (A) and *Ovgp1*-KO animals (B). Bars = 50 μm.

**Supplementary Table S1** Fertility of female individuals after ovarian transplantation.

* After the ovaries were implanted, they were mated with male individuals to see if they could produce litters.

** In 11 deliveries obtained after ovarian transplantation, the genotype of the fetus could not be confirmed in 5 cases due to maternal cannibalism immediately after delivery. The remaining 6 deliveries yielded a total of 29 fetuses (mean litter size = 4.83), 19 of which were *Ovgp1*^-/-^.

**Supplementary Table S2** List of proteins whose expression was changed after ovulation between WT and *Ovgp1*-KO hamsters oviducts. Of the 3573 proteins identified, 3/19 were significantly up/down regulated, respectively (these Volcano plots are shown in Figure 5).

## References

Araki, Y. (2022) Embryos, cancers, and parasites: potential applications to the study of reproductive biology in view of their similarity as biological phenomena. Reprod. Med. Biol. 21, e12447. doi: 10.1002/rmb2.12447

Araki, Y., Kurata, S., Oikawa, T., Yamashita, T., Hiroi, M., Naiki, M. and Sendo, F. (1987) A monoclonal antibody reacting with the zona pellucida of the oviductal egg but not with that of the ovarian egg of the golden hamster. J. Reprod. Immunol. 11, 193–208. doi: 10.1016/0165-0378(87)90057-x

Araki, Y., Orgebin-Crist, M.C. and Tulsiani, D.R. (1992) Qualitative characterization of oligosaccharide chains present on the rat zona pellucida glycoconjugates. Biol. Reprod. 46, 912–919. doi: 10.1095/biolreprod46.5.912

Araki, Y., Nohara, M., Yoshida-Komiya, H., Kuramochi, T., Ito, M., Hoshi, H., Shinkai, Y. and Sendai, Y. (2003) Effect of a null mutation of the oviduct-specific glycoprotein gene on mouse fertilization. Biochem. J. 374, 551–557. doi: 10.1042/BJ20030466.

Araki, Y. and Yoshida-Komiya, H. (1998) Mammalian oviduct-specific glycoprotein: Characterization and potential role in fertilization process. J. Reprod. Dev. 44, 215–228. doi:10.1262/jrd.44.215

Araki, Y., Yoshitake, H., Yamatoya, K. and Fujiwara, H. (2021) An overview of sex and reproductive immunity from an evolutionary/anthropological perspective. Immunol. Med. 44, 152–158. doi:10.1080/25785826.2020.1831219

Avilés, M., Gutiérrez-Adán, A. and Coy, P. (2010) Oviductal secretions: will they be key factors for the future ARTs? Mol. Hum. Reprod. 16, 896–906. doi: 10.1093/molehr/gaq056

Buhi, W. (2002) Characterization and biological roles of oviduct-specific, oestrogen-dependent glycoprotein. Reproduction 123, 355–362. doi:10.1530/rep.0.1230355

Edwards, R.G., Bavister, B.D. and Steptoe, P.C. (1969) Early stages of fertilization in vitro of human oocytes matured in vitro. Nature 221, 632–635. doi:10.1038/221632a0

Gardner, D.K., Lane, M., Calderon, I. and Leeton, J. (1996) Environment of the pre-implantation human embryo in vivo: Metabolite analysis of oviduct and uterine fluids and Metabolite analysis of oviduct and uterine fluids and metabolism of cumulus cells. Fertil. Steril. 65, 349–353. doi: 10.1016/s0015-0282(16)58097-2

González-Brusi, L., Algarra, B., Moros-Nicolás, C., Izquierdo-Rico, M.J., Avilés, M. and Jiménez-Movilla, M. (2020) A Comparative View on the Oviductal Environment during the Periconception Period. Biomolecules 10, 1690. doi: 10.3390/biom10121690.

Han, L., Monné, M., Okumura, H., Schwend, T., Cherry, A.L., Flot, D., Matsuda, T. and Jovine, L. (2010) Insights into egg coat assembly and egg-sperm interaction from the X-ray structure of full-length ZP3. Cell 143, 404–415. doi: 10.1016/j.cell.2010.09.041.

Hirose, M. and Ogura A. (2019) The golden (Syrian) hamster as a model for the study of reproductive biology: Past, present, and future. Reprod. Med. Biol. 18, 34–39. doi:10.1002/rmb2.12241

Hirose, M., Honda, A., Fulka, H., Tamura-Nakano, M., Matoba, S., Tomishima, T., Mochida, K., Hasegawa, A., Nagashima, K., Inoue, K., et al. (2020) Acrosin is essential for sperm penetration through the zona pellucida in hamsters. Proc. Natl. Acad. Sci. USA. 117, 2513–2518. doi: 10.1073/pnas.1917595117 (Erratum in: (2020) Proc. Natl. Acad. Sci. USA. 117, 24601.)

Huang, H.L., Li, S.C. and Wu, J.F. (2019) A complex of novel protease inhibitor, ovostatin homolog, with its cognate proteases in immature mice uterine luminal fluid. Sci. Rep. 9:4973. https://doi.org/10.1038/s41598-019-41426-4

Kan, F.W., Roux, E. and Bleau, G. (1993) Immunolocalization of oviductin in endocytic compartments in the blastomeres of developing embryos in the golden hamster. Biol. Reprod. 48, 77–88. doi: 10.1095/biolreprod48.1.77

Kapur, R.P. and Johnson, L, V. (1985) An oviductal fluid glycoprotein associated with ovulated mouse ova and early embryos. Dev. Biol. 112, 89–93. doi: 10.1016/0012-1606(85)90122-8

Kapur, R.P. and Johnson, L, V. (1986) Selective sequestration of an oviductal fluid glycoprotein in the perivitelline space of mouse oocytes and embryos. J. Exp. Zool. 238, 249–260. doi: 10.1002/jez.1402380215

Kapur, R.P. and Johnson, L, V. (1988) Ultrastructural evidence that specialized regions of the murine oviduct contribute a glycoprotein to the extracellular matrix of mouse oocytes. Anat. Rec. 221, 720–729. doi: 10.1002/ar.1092210307

Miyoshi, I., Takahashi, K., Kon, Y., Okamura, T., Mototani, Y., Araki, Y. and Kasai, N. (2002) Mouse transgenic for murine oviduct-specific glycoprotein promoter-driven simian virus 40 large T-antigen: tumor formation and its hormonal regulation. Mol. Reprod. Dev. 63, 168–176. doi: 10.1002/mrd.10175

Moros-Nicolás, C., Chevret, P., Jiménez-Movilla, M., Algarra, B., Cots-Rodríguez, P., González-Brusi, L., Avilés, M. and Izquierdo-Rico, M.J. (2021) New Insights into the Mammalian Egg Zona Pellucida. Int. J. Mol. Sci. 22, 3276. doi: 10.3390/ijms22063276

Oda-Sakurai R., Yoshitake, H., Miura, Y., Kazuno, S., Ueno, T., Hasegawa, A., Yamatoya, K., Takamori, K., Itakura, A., Fujiwara, H., et al. (2019) NUP62: the target of an anti-sperm auto-monoclonal antibody during testicular development. Reproduction 158, 503–516. doi: 10.1530/REP-19-0333

O’Day-Bowman, M.B, Mavrogianis, P.A., Minshall, R.D. and Verhage, H.G. (2002) In vivo versus in vitro oviductal glycoprotein (OGP) association with the zona pellucida (ZP) in the hamster and baboon. Mol. Reprod. Dev. 62, 248–256. doi: 10.1002/mrd.10091

Oikawa, T., Sendai, Y., Kurata, S. and Yanagimachi R. (1988) A glycoprotein of oviductal origin alters biochemical properties of the zona pellucida of hamster egg. Gamete Res. 19, 113–122. doi: 10.1002/mrd.1120190202

Quinn, P., Kerin, J.F. and Warnes, G.M. (1985a) Improved pregnancy rate in human in vitro fertilization with the use of a medium based on the composition of human tubal fluid. Fertil. Steril. 44, 493–498. doi: 10.1016/s0015-0282(16)48918-1

Quinn, P., Warnes, G.M., Kerin, J.F. and Kirby, C. (1985b) Culture factors affecting the success rate of in vitro fertilization and embryo transfer. Ann. N. Y. Acad. Sci. 442, 195–204. doi:10.1111/j.1749-6632.1985.tb37520.x

Robitaille, G., St-Jacques, S., Potier, M. and Bleau, G. (1988) Characterization of an oviductal glycoprotein associated with the ovulated hamster oocyte. Biol. Reprod. 38, 687–694. doi: 10.1095/biolreprod38.3.687. PMID: 2454136.

Roux, E., Bleau, G. and Kan, F.W. (1997) Fate of hamster oviductin in the oviduct and uterus during early gestation. Mol. Reprod. Dev. 46:306–317. doi: 10.1002/(SICI)1098-2795(199703)46:3<306::AID-MRD9>3.0.CO;2-T

Sakai, Y., Araki, Y., Yamashita, T., Kurata, S., Oikawa, T., Hiroi, M. and Sendo F. (1988) Inhibition of in vitro fertilization by a monoclonal antibody reacting with the zona pellucida of the oviductal egg but not with that of the ovarian egg of the golden hamster. J. Reprod. Immunol. 14, 177–189. doi: 10.1016/0165-0378(88)90068-x

Schini, S.A. and Bavister, B.D. (1988) Two-cell block to development of cultured hamster embryos is caused by phosphate and glucose. Biol. Reprod. 39, 1183–1192. doi: 10.1095/biolreprod39.5.1183

Sendai, Y., Komiya, H., Suzuki, K., Onuma, T., Kikuchi, M., Hoshi, H. and Araki Y. (1995) Molecular cloning and characterization of a mouse oviduct-specific glycoprotein. Biol. Reprod. 53, 285–294. doi: 10.1095/biolreprod53.2.285

Seshagiri, P.B. and Vani, V. (2019) Enabling Hamster Embryo Culture System: Development of Preimplantation Embryos. Methods Mol. Biol. 2006, 45–61. doi: 10.1007/978-1-4939-9566-0_4.

Takahashi, K., Sendai, Y., Matsuda, Y., Hoshi, H., Hiroi, M. and Araki Y. (2000) Mouse oviduct-specific glycoprotein gene: genomic organization and structure of the 5’-flanking regulatory region. Biol. Reprod. 62, 217–226. doi: 10.1095/biolreprod62.2.217

Takahashi, E,. Miyoshi, I. and Nagasu, T. Rescue of a transgenic mouse line by transplantation of a frozen-thawed ovary obtained postmortem. Contemp. Top. Lab. Anim. Sci. 40, 28–31. PMID: 11451392

Rankin, T., Familari, M., Lee, E., Ginsberg, A., Dwyer, N., Blanchette-Mackie, J., Drago, J., Westphal, H. and Dean J. (1996) Mice homozygous for an insertional mutation in the Zp3 gene lack a zona pellucida and are infertile. Development 122, 2903–2910. doi: 10.1242/dev.122.9.2903

Tulsiani, D.R, Yoshida-Komiya, H. and Araki, Y. Mammalian fertilization: a carbohydrate-mediated event. Biol. Reprod. 57, 487–494. doi: 10.1095/biolreprod57.3.487

Tumova, L,. Zigo, M., Sutovsky, P., Sedmikova, M. and Postlerova, P. (2021) Ligands and Receptors Involved in the Sperm-Zona Pellucida Interactions in Mammals. Cells 10, 133. doi: 10.3390/cells10010133.

Yanagimachi, R. and Chang, M.C. (1963) Fertilization of Hamster Eggs in vitro. Nature 200, 281–282. doi: 10.1038/200281b0

Yun, J.S., Li, Y.S., Wight, D.C., Portanova, R., Selden, R.F. and Wagner, T.E. (1990) The human growth hormone transgene: expression in hemizygous and homozygous mice. Proc. Soc. Exp. Biol. Med. 194, 308–313. doi: 10.3181/00379727-194-43096.

Yoshitake, H., Hashii, N., Kawasaki, N., Endo, S., Takamori, K., Hasegawa, A., Fujiwara, H. and Araki, Y. (2015) Chemical characterization of N-Linked oligosaccharide as the antigen epitope recognized by an anti-sperm auto-monoclonal antibody, Ts4. PLoS One 10, e0133784. doi: 10.1371/journal.pone.0133784

Zhao, Y., Vanderkooi, S. and Kan, F.W.K. (2022) The role of oviduct-specific glycoprotein (OVGP1) in modulating biological functions of gametes and embryos. Histochem. Cell Biol. 157, 371–388. doi: 10.1007/s00418-021-02065-x

